# Mice adaptively generate choice variability in a deterministic task

**DOI:** 10.1101/527580

**Authors:** Marwen Belkaid, Elise Bousseyrol, Romain Durand-de Cuttoli, Malou Dongelmans, Etienne K. Duranté, Tarek Ahmed Yahia, Steve Didienne, Bernadette Hanesse, Maxime Come, Alex Mourot, Jérémie Naudé, Olivier Sigaud, Philippe Faure

**Affiliations:** Sorbonne Université, CNRS, Institut des Systèmes Intelligents et de Robotique (ISIR), 75005 Paris, France; Sorbonne Université, INSERM, CNRS, Neuroscience Paris Seine - Institut de Biologie Paris Seine (NPS - IBPS), 75005 Paris, France

**Author notes:** correspondence should be addressed to P.F. Equally contributing senior authors (Olivier Sigaud, Philippe Faure).

## Abstract

Can decisions be made solely by chance? To investigate this question, we designed a deterministic setting in which mice are rewarded for non-repetitive choice sequences, and modeled the experiment using reinforcement learning. We found that mice progressively increased their choice variability using a memory-free, pseudo-random selection, rather than by learning complex sequences. Our results demonstrate that a decision-making process can self-generate variability and randomness even when the rules governing reward delivery are neither stochastic nor volatile.

## Main

Principles governing random behaviors are still poorly understood, despite well-known ecological examples ranging from vocal and motor babbling in trial-and-error learning ^1,2^ to unpredictable behavior in competitive setups (e.g preys-versus-predators ^3^ or humans competitive games ^4^). Dominant theories of behavior and notably reinforcement learning (RL) rely on exploitation, namely the act of repeating previously-rewarded actions ^5,6^. In this context, choice variability is associated with exploration of environmental contingencies. “Directed” exploration aims at gathering information about environmental contingencies ^7,8^, whereas random exploration introduces variability regardless of the contingencies ^9,10^. Studies have shown that animals are able to produce variable, unpredictable choices ^11,12^, especially when the reward delivery rule changes ^13,14^, is stochastic ^9,15,16^ or is based on predictions about their decisions ^17,18^. However, even approaches based on the prediction of the animal behavior ^17,18^ keep the possibility to distribute reward stochastically - for example if no systematic bias in the animal’s choice behavior has been found ^18,19^. Thus, because of the systematic use of volatile or probabilistic contingencies, it has remained difficult to experimentally isolate variability generation from environmental conditions. To test the hypothesis that animals can adaptively adjust the randomness of their behavior, we implemented a task where the reward delivery rule is deterministic, predetermined and identical for all animals, but where a purely random choices strategy is successful.

Mice were trained to perform a sequence of binary choices in an open-field where three target locations were explicitly associated with rewards delivered through intra-cranial self-stimulation (ICSS) in the medial forebrain bundle. Importantly, mice could not receive two consecutive ICSS at the same location. Thus, they had to perform a sequence of choices ^15^ and at each location to choose the next target amongst the two remaining alternatives (**Fig 1A**). In the training phase, all targets had a 100% probability of reward. We observed that after learning, mice alternated between rewarding locations following a stereotypical circular scheme interspersed with occasional changes in direction, referred to as U-turn (**Fig 1B**). Once learning was stabilized, we switched to the complexity condition, in which reward delivery was non-stochastic and depended on sequence variability. More precisely, we calculated the Lempel-Ziv (LZ) complexity ^20^ of choice subsequences of size 10 (9 past choices + next choice) at each trial. Animals were rewarded when they chose the one target (out of the two options) associated with the highest complexity (given the previous nine choices). Despite its difficulty, this task is fully deterministic. Indeed, mice were asked to move along a tree of binary choices (see **Fig 1A**) where some paths ensured 100% rewards. Whether each node was rewarded or not was pre-determined in advance. Thus, choice variability could not be imputed to the inherent stochasticity of the outcomes. For each trial, if choosing randomly, the animal had either 100% or 50% chance of being rewarded depending on whether the two subsequences of size 10 (= 9 past choices + 1 choice out of 2 options) had equal or unequal complexities. Another way to describe the task is thus to consider all possible situations, not as sequential decisions made by the animal during the task but as the set of all possible subsequences of size 10 of which the algorithm may evaluate the complexity. From this perspective, there is an overall 75% probability of being rewarded if subsequences are sampled uniformly (**Fig 1A**). To summarize, theoretically, while a correct estimation of the complexity of the sequence leads to a success rate of 100%, a pure random selection leads to 75% of success, and a repetitive sequence (e.g. A,B,C,A,B,C,…) grants no reward.

**Figure 1:**
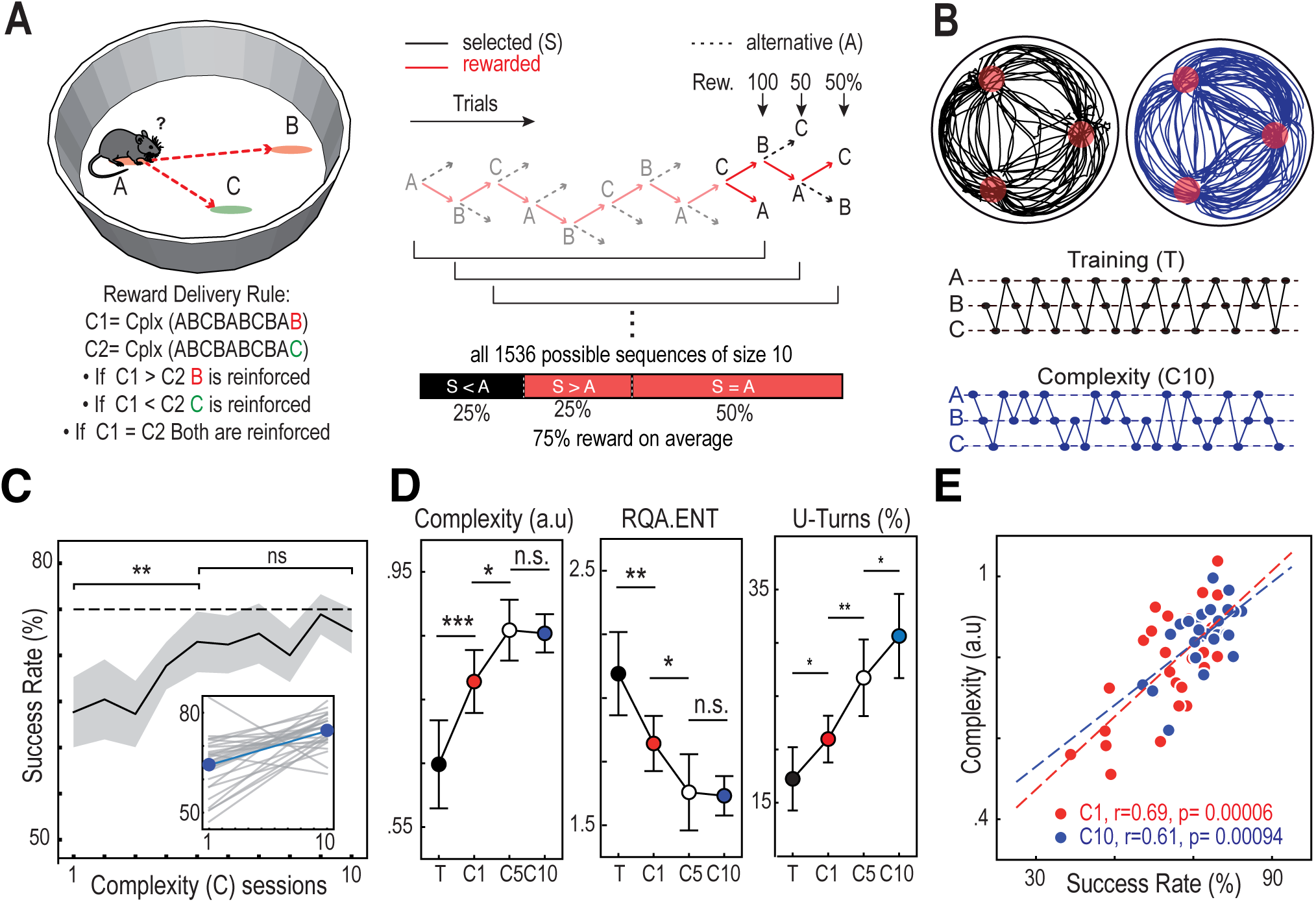
Mice generate unpredictable decisions. **A)** *Left:* Task setting. Mice were trained to perform a sequence of binary choices between the two out of three target locations (A, B and C) associated with ICSS rewards. Under the complexity condition, for each trial, the grammatical complexity of two subsequences of size 10 (last 9 choices concatenated with the two options) were compared. Animals were rewarded when they chose the target giving the highest complexity. *Right:* Tree structure of the task and reward distribution. Giving that the reward delivery is deterministic, the task can be seen as a decision tree in which some paths ensure 100% rewards. From a local perspective, for each trial, the animal has either 100% or 50% chance of reward; resp. if the evaluated subsequences of size 10 have equal or unequal complexities. Considering all these possible sequences, 75% of the trials would be rewarded if animals were to choose randomly. **B)** Typical trajectories before and after switching to the complexity condition. In the training phase (before complexity), mice alternated between the equally rewarded targets following a stereotypical circular scheme (e.g. ABCABCABC…). At the end of the complexity condition, choice sequence became more variable. **C)** Increase of the success rate over sessions in the complexity setting. Mice improved their performance in the first sessions (c01 versus c05, T = 223.5, p = 0.015, Wilcoxon test) then reached a plateau (c05 versus c10, t(25) = −0.43, p = 0.670, paired t-test) close to the theoretical 75% success rate of random selection (c10, t(25) = −1.87, p = 0.073, single sample t-test). The shaded area represents a 95% confidence interval. *Inset*, grey lines represent linear regressions of individual mice performance increase for individual mice and the blue line represents the average progress. **D)** Increase of the behavior complexity over sessions. *Left*: the *NLZcomp* measure of complexity increased in the beginning (training versus c01, T = 52, p = 0.0009, Wilcoxon test, c01 versus c05, t(26) = −2.67, p = 0.012, paired t-test) before reaching a plateau (c05 versus c10, T = 171, p = 0.909, Wilcoxon test). The average complexity reached by animal is lower than 1 (c10, t(25) = −9.34, p = 10-9, single sample t-test), which corresponds to the complexity of random sequences. *Middle:* the RQA *ENT* entropy-based measure of complexity decreased over sessions (training versus c01, t(26) = 2.81, p = 0.009, paired t-test, c01 versus c05, T = 92, p = 0.019, Wilcoxon test, c05 versus c10, T = 116, p = 0.13, Wilcoxon test). *Right*: The rate of U-turns increased over sessions (training versus c01, t(26) = −2.21, p = 0.036, c01versus c05, t(26) = −3.07, p = 0.004, paired t -test, c05 versus c10, T = 75, p = 0.010, Wilcoxon test). Error bars represent 95% confidence intervals. **E)** Correlation between individual success rate and complexity of mice sequences. Also noteworthy is the decrease in data dispersion in session c10 compared to c1. N = 27 in all sessions except c10 where N = 26. * p < 0.05, ** p < 0.01, *** p < 0.001. n.s., not significant at p > 0.05.

**Table 1:** Hyperparameter ranges used in random search in independent, session-by-session model fitting (*N*_*samples*_ = 6000) and in grid search in continuous, all-sessions model fitting.

| Hyperparameter random search ranges |  |  |  |
| --- | --- | --- | --- |
| Label | Range | Step | Description |
| $m$ | $[0, 9]$ | 1 | Memory size |
| $\tau$ | $[1, 20]$ | *continuous* | Softmax temperature |
| $\kappa$ | $[0, 1]$ | *continuous* | U-turn cost |

| Hyperparameter grid search ranges |  |  |  |
| --- | --- | --- | --- |
| Label | Range | Step | Description |
| $m$ | $[0, 9]$ | 1 | Memory size (no ambiguity) |
| | $[0, 7]$ | 1 | Memory size (low ambiguity) |
| | $[0, 5]$ | 1 | Memory size (medium ambiguity) |
| $\alpha$ | $2^{-i}, i \in [0, 10]$ | 1 | Learning rate |
| $\tau$ | $2^i, i \in [-4, 4]$ | 1 | Softmax temperature |
| $\kappa$ | $[0.5, 0.95]$ | 0.05 | U-turn cost |

We found that mice progressively increased the variability of their choice sequences (**Fig 1B**) and thus their success rate along sessions (**Fig 1C**). This increased variability in the generated sequences was demonstrated by an increase in the normalized LZ-complexity measure (hereafter *NLZcomp*) of the session sequences, a decrease in an entropy measure based on recurrence plot quantification and an increase in the percentage of U-turns (**Fig 1D**). Furthermore, in the last session, 65.5% of the sequences were not significantly different from surrogate sequences generated randomly (**Supp Fig 1A**). The success rate was correlated with the *NLZcomp* of the entire session of choice sequences (**Fig 1E**), suggesting that mice increased their reward through an increased variability in their choice. The increase in success rate was associated with an increase of the percentage of U-turns (**Fig 1D** left), yet mice performed a suboptimal U-turn rate of 30%, below the 50% U-turn rate ensuring 100% of rewards (**Supp Fig 1B**).

From a behavioral point of view, mice thus managed to increase their success rate in a highly demanding task. They did not achieve 100% success but reached performances that indicate a significant level of variability. Given that the task is fully deterministic, the most efficient strategy would be to learn and repeat one (or some) of the 10-choice long sequences that are always rewarded. This strategy ensures the highest success rate but incurs a tremendous memory cost. On the other hand, a purely random selection is another appealing strategy since it is less costly and leads to about 75% of reward. To differentiate between the two strategies and better understand the computational principles underlying variability generation in mice, we examined the ability of a classical RL algorithm to account for the mouse decision-making process under these conditions.

State-action values were learned using the Rescorla-Wagner rule ^21^ and action selection was based on a softmax policy ^5^ (**Fig 2A;** see ‘Methods’). By defining states as vectors including the history of previous locations instead of the current location alone, we were able to vary the memory size of simulated mice and to obtain different solutions from the model accordingly. We found that, with no memory (i.e. state = current location), the model learned equal values for both targets in almost all states (**Fig 2B**). In contrast, and in agreement with classical RL, with the history of the nine last choices stored in memory, the model favored the rewarded target in half of the situations by learning higher values (approximately 90 vs 10%) associated with rewarded sequences of choices (**Fig 2B**). This indicates that classical RL can find the optimal solution of the task if using a large memory. Furthermore, choosing randomly was dependent not only on the values associated with current choices, but also on the softmax temperature and the U-turn cost. The ratio between these two hyperparameter controls the level of randomness in action selection (see ‘Methods’). Intuitively, a high level of randomness leads to high choice variability and sequence complexity. But interestingly, the randomness hyperparameter had opposite effects on the model behavior with small and large memory sizes. While increasing the temperature always increased the complexity of choice sequences, it increased the success rate for small memory sizes but decreased it for larger memories (**Fig 2C**). A boundary between the two regimes was found between memory sizes of 3 and 4.

**Figure 2:**
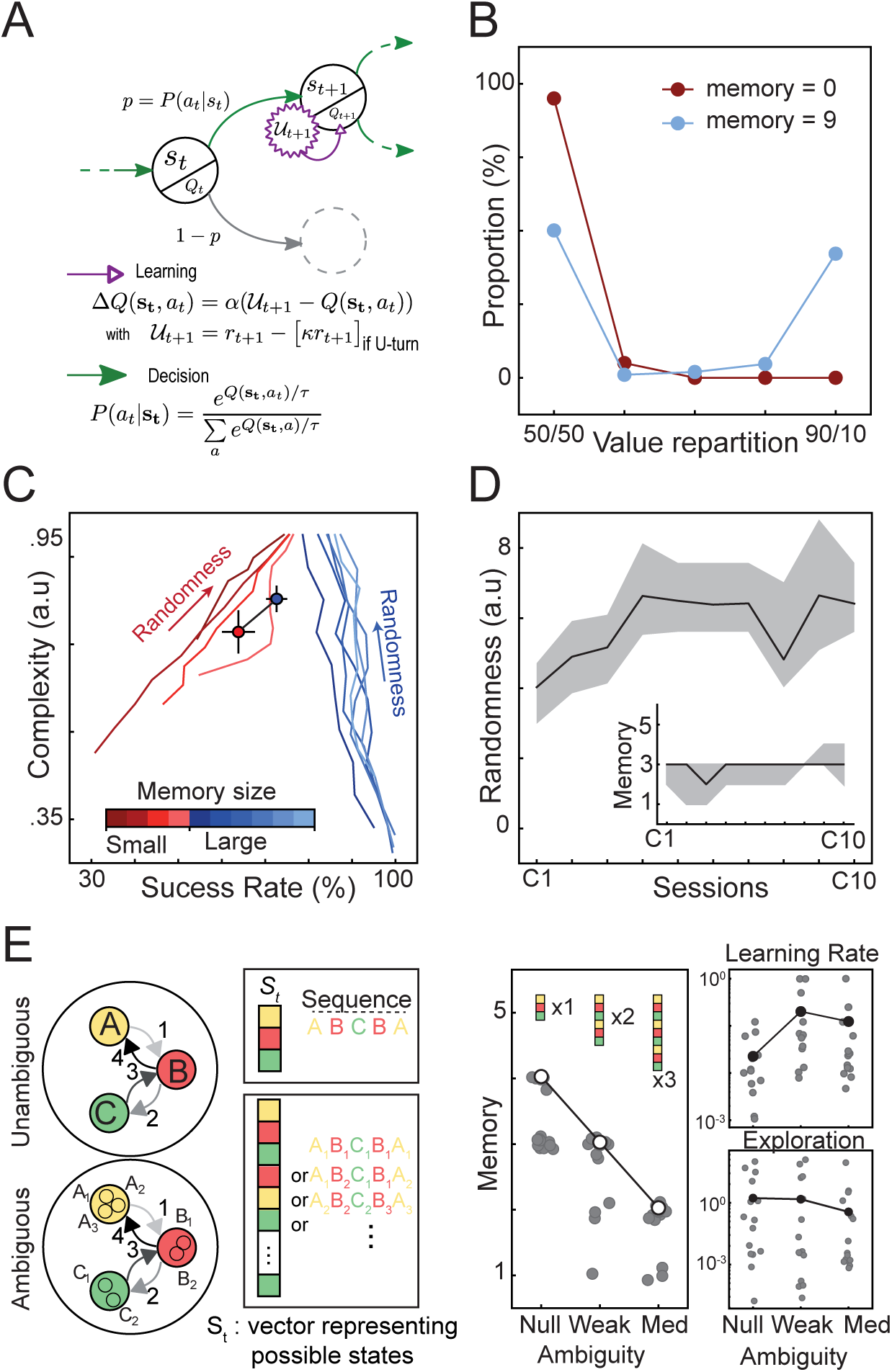
Computational modeling suggests a memory-free pseudo-random selection behind the mice generation of variability. **A)** Schematic illustration of the computational model fitted to mouse behavior. As in classical reinforcement learning, state-action values were learned using the Rescorla-Wagner rule and action selection was based on a softmax policy. Two adaptations were applied: i) rewards were discounted by a U-turn cost 𝒳 in the utility function in order to reproduce mouse circular trajectories in the training phase (see ‘Methods’) ii) states were represented as vectors in order to simulate mouse memory of previous choices. **B)** Repartition of the values learned by the model with memory size equal to 0 or 9. With a large memory, the model is able to favor the rewarded target (approx. 90/10 repartition) in the situations where the two options are not equally rewarded (see Fig 1A). With no memory, both actions have about the same value (approx. 50/50 repartition) in almost 100% of the states. **C)** Influence of increased randomness on success rate and complexity for various memory sizes. Each line describes the trajectory followed by a model with a certain memory size (see color scale) when going from a low to high level of randomness. The randomness hyperparameter is defined as **τ** / *κ*. Red and blue dots represent experimental data of mice in the last training and complexity sessions respectively. Error bars represent 95% confidence intervals. **D)** Model fitting results. With an increase of exploration and a small memory, the model fits the increase in mice performance. The shaded areas represent values of the 15 best parameter sets. Dark lines represent the average randomness value (continuous values) and the best fitting memory size (discrete values) respectively. **E)** Schematic of the difference between ambiguous and unambiguous state representations and simulation results. Top-Left: The main simulations rely on an unambiguous representation of states in which each choice sequence is represented by one perfectly recognized code. Bottom-Left: With more ambiguous states, the same sequence can be encoded by a variety of representations. Right: With higher representation ambiguity, the model better fits mouse performance with a smaller memory (null, weak and medium ambiguity, H = 27.21, p = 10-6, Kruskal-Wallis test, null versus weak, U = 136, p = 0.006, weak versus med, U = 139, p = 0.002, Mann-Whitney test) and with a higher learning rate (null, weak and medium ambiguity, H = 7.61, p = 0.022, Kruskal-Wallis test, null versus weak, U = 45.5, p = 0.016, null versus med, U = 54, p = 0.026, weak versus med, U = 101, p = 0.63, Mann-Whitney test) but a similar exploration rate (null, weak and medium ambiguity, H = 3.64, p = 0.267, Kruskal-Wallis test). Grey dots represent the 15 best fitting parameter sets. White dots represent the best fit in case of a discrete variable (memory) while black dots represent the average in case of continuous variables (temperature and learning rate). N = 15.

Upon optimization of the model to fit mouse behavior, we found that their performance improvement over sessions was best accounted for by an increase of choice randomness using a small memory (**Fig 2D**). This model captured mouse learning better than when using fixed parameters throughout sessions (Bayes factor = 3.46; see ‘Methods’, and **Supp Fig 2D and E**). The model with a memory of size 3 best reproduced mouse behavior (**Fig 2D**), but only slightly better than versions with smaller memories (**Supp Fig 2C**). From a computational perspective, one possible explanation for the fact that although theoretically sufficient, a memory of size 1 fits less than size 3, is that state representation is overly simplified in the model. Accordingly, altering the model’s state representation to make it more realistic should reduce the size of the memory needed to reproduce mice performances. To test this hypothesis, we used a variant of the model in which we manipulated state representation ambiguity: each of the locations *{A, B, C}* could be represented by *n ≥1* states, with *n = 1* corresponding to unambiguous states (see ‘Methods’, and **Fig 2E**). As expected, the model fitted better with a smaller memory as representation ambiguity was increased (**Fig 2E**). We also found that the best fitting learning rate was higher with ambiguous representations while the randomness factor remained unchanged regardless of ambiguity level (**Fig 2E**). This corroborates that the use of additional memory capacity by the model is due to the model’s own limitations rather than an actual need to memorize previous choices. Hence, this computational analysis overall suggests that mice adapted the randomness parameter of their decision-making system to achieve more variability over sessions rather than remembered rewarded choice sequences. This conclusion was further reinforced by a series of behavioral arguments detailed below supporting the lack of memorization of choice history in their strategy.

We first looked for evidence of repeated choice patterns in mouse sequences using a Markov chain analysis (see ‘Methods’). We found that the behavior at the end of the complexity condition was Markovian (**Fig 3A**). In other words, the information about the immediately preceding transition (i.e. to the left or to the right) was necessary to determine the following one (e.g. p(L) ≠ P(L|L)) but looking two steps back was not informative on future decisions (e.g. p(L|LL) ≈ P(L|L)). The analysis of the distribution of subsequence of length 10 (see Methods) provides an additional evidence of the lack of structure in the animals’ choice sequence. Indeed, while at the end of the training, mice exhibit a peaky distribution with a strong preference for the highly repetitive circular patterns, the distribution was dramatically flattened under the complexity condition (**Fig 3B**). Furthermore, we tested whether mice use of a win-stay-lose-switch strategy ^17^. Indeed, mice could have used this heuristic strategy when first confronted with the complexity condition after a training phase in which all targets were systematically rewarded. Changing directions in the absence of reward could have introduced enough variability in the animals’ sequence to improve their success rate. Yet, we found that being rewarded (or not) had no effect on the next transition, neither at the beginning nor the end of the complexity condition (**Fig 3C**); thus eliminating another potential form of structure in mice behavior under the complexity rule.

**Figure 3:**
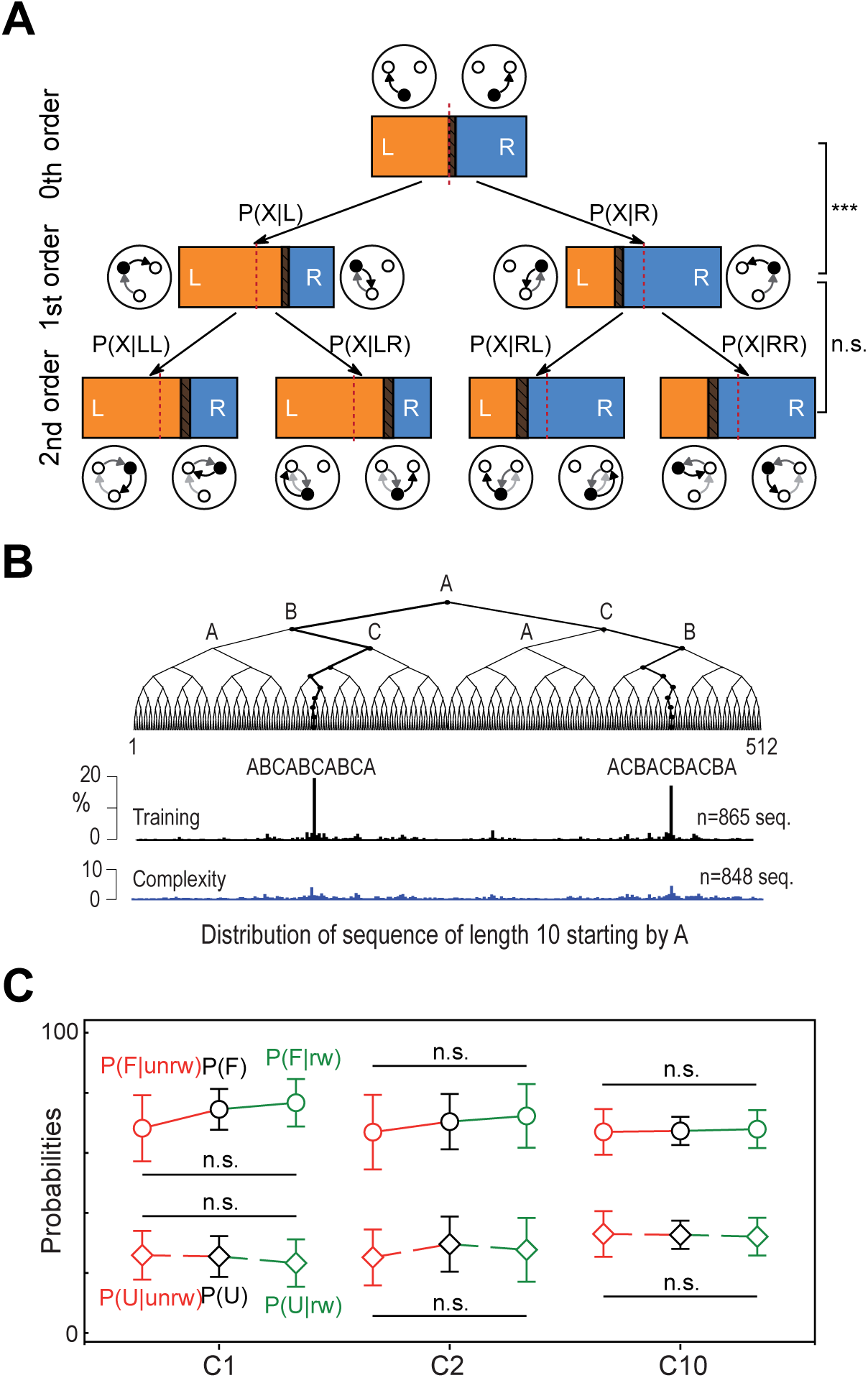
Behavioral evidence of the absence of memorization in mouse choices. **A)** Tree representation of the Markovian structure of mouse behavior in session c10 (N = 26). In the expression of probabilities, P(X) refers to P(L) or P(R), whose repartition is illustrated in the horizontal bars (respectively in orange and blue). Dashed areas inside the bars represent overlapping 95% confidence intervals. The probability of a transition (i.e. to the left or to the right) is different from the probability of the same transition given the previous one (p(L) versus P(L|L), t(25) = −7.86, p = 3.10-8, p(L) versus P(L|R), t(25) = 7.57, p = 6.10-8, p(R) versus P(R|R), t(25) = −7.57, p = 6.10-8, p(R) versus P(R|L), t(25) = 7.86, p = 3.10-8, paired t-test). However, the probability given two previous transitions is not different from the latter (p(L|L) versus P(L|LL), t(25) = 1.36, p = 0.183, p(L|L) versus P(L|LR), t(25) = −1.66, p = 0.108, p(L|R) versus P(L|RL), t(25) = −0.05, p = 0.960, p(L|R) versus P(L|RR), t(25) = −0.17, p = 0.860, p(R|R) versus P(R|RR), t(25) = 0.17, p = 0.860, p(R|R) versus P(R|RL), t(25) = 0.05, p = 0.960, p(R|L) versus P(R|LR), t(25) = 1.66, p = 0.108, p(R|L) versus P(R|LL), t(25) = −1.36, p = 0.183, paired t-test). **B)** Distribution of subsequences of length 10. All subsequences starting by A were extracted from the choice sequences of mice in the last sessions of the training condition and the complexity condition. The histograms represent their distribution following the tree structure to highlight subsequences similarities. Under the training condition, mice show a preference for the two most repetitive patterns and their variants. In contrast, the distribution is markedly flattened under the complexity condition demonstrating that mice behavior is much less structured in this setting. **C)** Absence of influence of rewards on mice decisions. P(F) and P(U) respectively refer to the probabilities of going forward (e.g. A→B→C) and making a U-turn (e.g. A→B→A). These probabilities were not different from the conditional probabilities given that the previous choice was rewarded or not (c01, P(F), P(F|rw) and P(F|unrw), H = 2.93, p = 0.230, P(U), P(U|rw) and P(U|unrw), H = 1.09, p = 0.579, c02, P(F), P(F|rw) and P(F|unrw), H = 1.08, p = 0.581, P(U), P(U|rw) and P(U|unrw), H = 0.82, p = 0.661, c10, P(F), P(F|rw) and P(F|unrw), H = 0.50, p = 0.778, P(U), P(U|rw) and P(U|unrw), H = 0.50, p = 0.778, Kruskal-Wallis test). This means that the change in mice behavior under the complexity condition was not stereotypically driven by the outcome of their choices (e.g. ‘u-turn if not rewarded’). Error bars in B represent 95% confidence intervals. N = 27 in c01 and c02 and N = 26 in c10. Error bars represent 95% confidence intervals. * p < 0.05, ** p < 0.01, *** p <0.001. n.s., not significant at p > 0.05.

To further support the notion that mice did not actually memorize rewarded sequences to solve the task, we finally performed a series of experiments to compare the animals’ behavior under the complexity rule and under a probabilistic rule in which all targets were rewarded with a 75% probability (the same frequency reached at the end of the complexity condition). We first analyzed mice behavior when the complexity condition was followed by the probabilistic condition (Group 1 **Fig 4A**). We hypothesized that, if animals choose randomly at each node in the complexity setting (and thus do not memorize and repeat specific sequences), they would not detect the change of the reward distribution rule when switching to the probabilistic setting. In agreement with our assumption, we observed that as we switched to the probabilistic condition, animals did not modify their behavior although the optimal strategy would have been to avoid U-turns, as observed in the 100% reward setup used for training (**Fig 4B** and **Supp Fig 3A**). Hence, after the complexity setting, mice were likely stuck in a “random” mode given that the global statistics of the reward delivery were conserved. In contrast, when mice were exposed to the probabilistic distribution of reward right after the training session (Group 2 **Fig 4A**), they slightly changed their behavior but mostly stayed in a circular pattern with few U-turns and low sequence complexity (**Fig 4B** and **Supp Fig 3A**). Thus, animals from Group 2 exhibited lower sequence complexity and U-turn rate in the probabilistic condition than animals from Group 1, whether in the complexity or the probabilistic condition (**Fig 4C**). The distribution of patterns of length 10 in the sequences performed by animals from Group 2 during the last probabilistic session shows a preference for repetitive circular patterns that is very similar to that observed at the end of the training; contrasting with the sequences performed by animals from Group 1 (**Fig 4D**). A larger portion of sequences performed by animals from Group 1 were not significantly different from surrogate sequences generated randomly in comparison with animals from Group 2 (**Supp Fig 3B**). Last, if the sequences performed by mice from Group 2 were executed under the complexity rule, these animals would have obtained significantly lower success rate that animals from Group 1 in the complexity condition (**Supp Fig 3C**). In summary, mice behavior under the probabilistic condition changed significantly depending on the preceding condition and the strategy that the animal was adopting. This further supports our initial claim that stochastic experimental setups made it difficult to unravel the mechanisms underlying random behavior generation. On the other hand, the complexity rule used in our experiments make it possible to categorize animals’ behavior into one of three possible strategies (i.e. repetitive, random or optimal). Overall, our results indicate no evidence of sequence memorization nor any behavioral pattern that might have been used by mice as a heuristic to solve the complex task.

**Figure 4:**
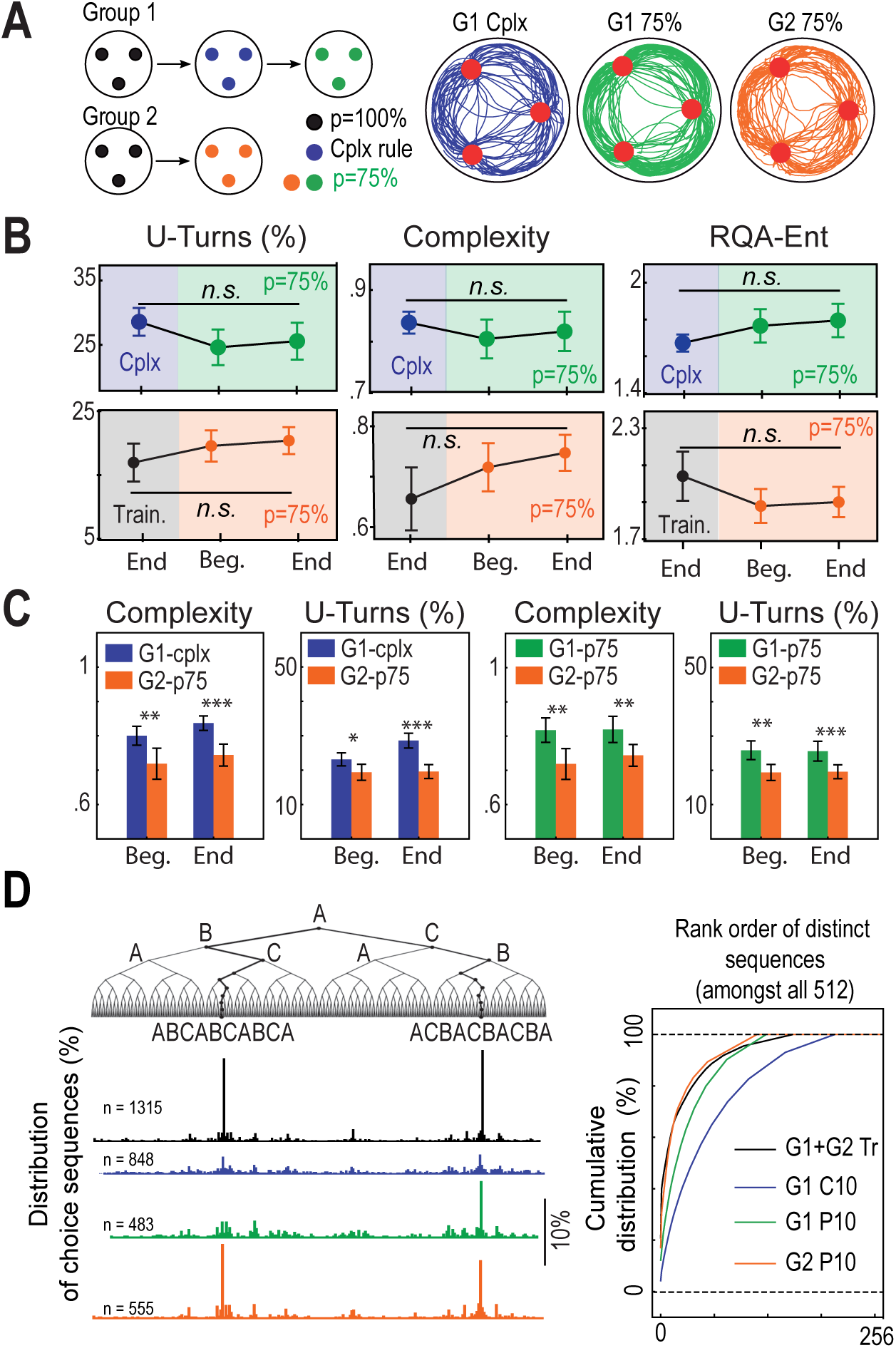
Comparison of mice behavior under the complexity condition and a probabilistic condition. **A)** *Left:* Schematic illustration of the sequence of conditions in two new experiments. For a first group of mice (G1), the complexity condition was followed by a probabilistic condition. For a second group (G2), the probabilistic condition was experienced right after the training phase. Under the probabilistic condition, choices were rewarded with the same frequency reached at the end of the competitor condition but in a stochastic way. *Right*: Typical trajectories under the complexity condition and the probabilistic condition. **B)** Analysis of the behavior of G1 and G2 mice in the probabilistic setting compared to the end of the preceding condition (resp. complexity and training condition). For G1 animals, the U-turn rate, *NLZcomp* complexity measure and RQA *ENT* measure remain unchanged (pooled ‘end vc’, ‘beg. p75’ and ‘end p75’, U-turn rate, H = 4.22, p = 0.120, Complexity, H = 0.90, p = 0.637, RQA ENT, H = 4.57, p = 0.101, Kruskal-Wallis test). For G2 animals as well, the U-turn rate, *NLZcomp* complexity measure and RQA *ENT* measure remain unchanged (pooled ‘end tr’, ‘beg. p75’ and ‘end p75’, U-turn rate, H = 5.68, p = 0.058, Complexity, H = 4.10, p = 0.128, RQA ENT, H = 2.66, p = 0.073, Kruskal-Wallis test). **C)** Comparison of G2 behavior in the probabilistic setting with G1 behavior under the complexity (Left) and the probabilistic (Right) conditions. G1 mice exhibit higher sequence complexity and U-turn rate than G2 under both the complexity condition (G1-cplx versus G2-p75, Complexity, pooled ‘beg’, t(136) = 2.99, p = 0.003, pooled ‘end’, t(136) = 4.72, p = 7.10^−6^, Welch t-test, U-turn, pooled ‘beg’, U = 2866.5, p = 0.015, pooled ‘end’, U = 3493, p = 10^−7^, Mann-Whitney test) and the probabilistic condition (G1-p75 versus G2-p75, Complexity, pooled ‘beg’, U = 1375, p = 0.005, Mann-Whitney test, pooled ‘end’, t(91) = 2.92, p = 0.004, t-test, U-turn, pooled ‘beg’, U = 1478, p = 0.0003, pooled ‘end’, U = 1424, p = 0.001, Mann-Whitney test). N = 80 for G1-cplx, N = 36 for G1-p75, N = 54 for G2-p75. **D)** *Left*: Distribution of subsequences of length 10 performed by G1 and G2 animals in the last sessions of the training, the complexity (same as in Fig 3B) and the probabilistic conditions represented following the tree structure. Under the probabilistic condition, mice from Group 2 show a preference for the two most repetitive patterns and their variants, similarly to the training condition. Such structure is present in mice from Group 1 but in a lesser extent. *Right*: Cumulative distribution of ranked patterns of length 10. There is a marked difference between the sequences performed under the complexity rule in comparison to the training and the probabilistic condition. Error bars represent 95% confidence intervals. * p < 0.05, ** p < 0.01, *** p < 0.001. n.s., not significant at p > 0.05.

Whether and how the brain can generate random patterns has always been puzzling ^22^. In this study, we addressed two fundamental aspects in this matter: the implication of memory processes and the dependence upon external (environmental) factors. Regarding memory, one hypothesis holds that in human, the process of generating random patterns leverages memory ^23^, to ensure the equality of response usage for example ^24^, whereas a second hypothesis suggests that the lack of memory may help eliminate counterproductive biases ^25 26^. In our task, mice did not use their memory, thus suggesting that the brain is able to effectively achieve high variability by suppressing biases and structure, at least in some contexts. The second aspect is the degree of dependence upon external, environmental factors. Exploration and choice variability are generally studied by introducing stochasticity and/or volatility in environmental outcomes ^15-18^. However, such conditions make it difficult to interpret the animal’s strategy and to know whether the observed variability in the mouse choice is inherited or not from the statistics of the behavioral task. In this work, we took a step further toward understanding the processes underlying the generation of variability per se, independently from environmental conditions. Confronted with a deterministic task which yet favors complex choice sequences, mice avoided repetitions by engaging in a behavioral mode where decisions were random and independent from their reward history. Animals adaptively tuned their decision-making parameters to increase choice randomness, which suggests an internal process of randomness generation.

## Methods

### Animals

Male C57BL/6J (WT) mice obtained from Charles Rivers Laboratories France (L’Arbresle Cedex, France) were used. Mice arrived to the animal facility at 8 weeks of age, and were housed individually for at least 2 weeks before the electrode implantation. Behavioral tasks started one week after implantation to ensure full recovery. Since intracranial self-stimulation (ICSS) does not require food deprivation, all mice had ad libitum access to food and water except during behavioral sessions. The temperature (20-22 °C) and humidity was automatically controlled and a circadian light cycle of 12/12h light-dark cycle (lights on at 8:30 am) was maintained in the animal facility. All experiments were performed during the light cycle, between 09:00 a.m. and 5:00 p.m. Experiments were conducted at Sorbonne University, Paris, France, in accordance with the local regulations for animal experiments as well as the recommendations for animal experiments issued by the European Council (directives 219/1990 and 220/1990).

### ICSS

Mice were introduced into a stereotaxic frame and implanted unilaterally with bipolar stimulating electrodes for ICSS in the medial forebrain bundle (MFB, anteroposterior = 1.4 mm, mediolateral = ±1.2 mm, from the bregma, and dorsoventral = 4.8 mm from the dura). After recovery from surgery (1 week), the efficacy of electrical stimulation was verified in an open field with an explicit square target (side = 1 cm) at its center. Each time a mouse was detected in the area (D = 3 cm) of the target, a 200-ms train of twenty 0.5-ms biphasic square waves pulsed at 100 Hz was generated by a stimulator. Mice self-stimulating at least 50 times in a 5 minutes session were kept for the behavioral sessions. In the training condition, ICSS intensity was adjusted so that mice self-stimulated between 50 and 150 times per session at the end of the training (ninth and tenth session), then the current intensity was kept the same throughout the different settings.

### Complexity task

In the complexity condition, reward delivery was determined by an algorithm that estimated the grammatical complexity of animals’ choice sequences. More specifically, at a trial in which the animal was at the target location A and had to choose between B and C, we compared the LZ-complexity ^20^ of the subsequences comprised of the 9 past choices and B or C. Both choices were rewarded if those subsequences were of equal complexity. Otherwise, only the option making the subsequence of highest complexity was rewarded.

### Measures of choice variability

Two measures of complexity were used to analyze mouse behavior. First, the normalized LZ-complexity (referred to as *NLZcomp* or simply *complexity* throughout the paper) which corresponds to the LZ-complexity divided by the average LZ-complexity of 1000 sequences of the same length generated randomly (a surrogate) with the constraint that two consecutive characters could not be equal, as in the experimental setup. *NLZcomp* is small for highly repetitive sequence and is close to 1 for uncorrelated, random signals. Second, the entropy of the frequency distribution of the diagonal length (noted *ENT*), taken from recurrence quantification analysis (RQA). RQA is a series of methods in which the dynamics of complex systems are studied using recurrence plots (RP) ^27,28^ where diagonal lines illustrate recurrent patterns. Thus, the entropy of diagonal lines reflects the deterministic structure of the system and is smaller for uncorrelated, random signals. RQA was measured using the Recurrence-Plot Python module of the “pyunicorn.timeseries*”* package.

### Computational models

The task was represented as a Markov Decision Process (MDP) with three states *s ∈* {A, B, C} and three actions *a ∈* {GoToA, GoToB, GoToC}, respectively corresponding to the rewarded locations and the transitions between them. State-action values *Q(s,a)* were learned using the Rescorla-Wagner rule ^21^:

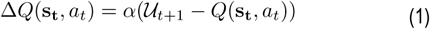

where ***s*** _***t***_ =[*s*_*t*_, *s*_*t-1*_, *…, s*_*t-m*_] is the current state which may include the memory of up to the m^th^ past location, a_t_ the current action, α the learning rate and *U* the utility function defined as follows:

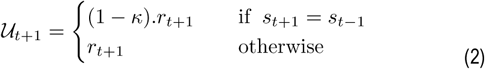

where *r* is the reward function and *κ* the U-turn cost parameter modeling the motor cost or any bias against the action leading the animal back to its previous location. The U-turn cost was necessary to reproduce mouse stereotypical trajectories at the end of the training phase (see **Supp Fig 2**).

Action selection was performed using a softmax policy, meaning that in state *s*_*t*_ the action *a*_*t*_ is selected with probability:

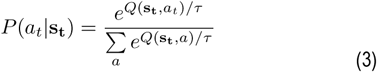

where **τ** is the temperature parameter. This parameter reduces the sensitivity to the difference in actions values thus increasing the amount of noise or randomness in decision-making. The U-turn cost *κ* has the opposite effect since it represents a behavioral bias and constrains choice randomness. We refer to the hyperparameter defined as *ρ*= ***τ*** / *κ* as the randomness parameter.

In the version referred to as *BasicRL* (see **Supp Fig 2**), we did not include any memory of previous locations nor any U-turn cost. In other words, *m=0* (i.e. ***s*** _***t***_ = [*s*_*t*_]) and *κ* = 0.

To manipulate state representation ambiguity (see **Fig 2**), each of the locations {A, B, C} could be represented by *n≥1* states. For simplicity, we used *n=*1, 2 and 3 for all locations for what we referred to as ‘null’, ‘low, and ‘med’ levels of ambiguity. This allowed us to present a proof of concept regarding the potential impact of using a perfect state representation in our model.

### Model Fitting

The main model-fitting results presented in this paper were obtained by fitting the behavior of the mice under training and complexity conditions session by session independently. This process aimed to determine which values of the two hyperparameters m and *ρ*= ***τ*** / *κ* make the model behave as mice in terms of success rate (i.e. percentage of rewarded actions) and complexity (i.e. variability of decisions). Our main goal was to decide between the two listed strategies that can solve the task: repeating rewarded sequences or choosing randomly. Therefore, we momentarily put aside the question of learning speed and only considered the model behavior after convergence. α was set to 0.1 in these simulations.

Hyperparameters were selected through random search ^29^ (see ranges listed in **Supplementary Table 1**). The model was run for 2.10^6^ iterations for each parameter set. The fitness score with respect to mice average data at each session was calculated as follows:

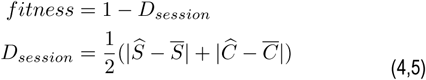

where 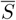 and 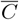 are the average success rate and complexity in mice respectively and *Ŝ* and *Ĉ* the model success rate and complexity -- all the four *∈* [0,1]. Simulations were long enough for the learning to converge. Thus, instead of multiple runs for each parameter set, which would have been computationally costly, *Ŝ* and *Ĉ* were averaged over the last 10 simulated sessions. We considered that 1 simulated session = 200 iterations, which is an upper bound for the number of trials performed by mice in one actual session.

Since mice were systematically rewarded during training, their success rate under this condition was not meaningful. Thus, to assess the ability of the model to reproduce stereotypically circular trajectories in the last training session, we replaced *Ŝ* and 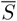 in Equation (5) by *Û* and *Ū* representing the average U-turn rates for mice and for the model respectively.

Additional simulations were conducted with two goals: 1) test whether one single parameter set could fit mice behavior without the need to change parameter values over sessions, 2) test the influence of state representation ambiguity on memory use in the computational model. Therefore, each simulation attempted to reproduce mice behavior from training to the complexity condition. Hence, the learning rate α was optimized in addition to the previously mentioned m and *ρ* = ***τ*** / *κ* hyperparameters (see ranges listed in **Supplementary Table 1**). Each parameter set was tested over 20 different runs. Each run is a simulation of 4000 iterations, which amounts to 10 training sessions and 10 complexity sessions since simulated sessions consist of 200 iterations. The fitness score was computed as the average score over the last training session and the 10 complexity sessions using Equations (4) and (5). Using a grid search ensured comparable values for different levels of ambiguity (‘null’, ‘low, and ‘med’; see previous section). Given the additional computational cost induced by higher ambiguity levels, we gradually decreased the upper bound of the memory size range in order to avoid long and useless computations in uninteresting regions of the search space.

### Markov chain analysis

Markov chain analysis allows to mathematically describe the dynamic behavior of the system, i.e. transitions from one state to another, in probabilistic terms. A process is a first-order Markov chain (or more simply *Markovian*) if the transition probability from state A to a state B depends only on the current state A and not on the previous ones. Put differently, the current state contains all the information that could influence the realization of the next state. A classical way to demonstrate that a process is Markovian is to show that the sequence cannot be described by a zeroth-order process, i.e. that P(B|A) ≠ P(B), and that the second-order probability is not required to describe the state transitions, i.e. that P(B|A) = P(B|AC).

In this paper, we analyzed the 0^th^, 1^st^ and 2^nd^ order probabilities in sequences performed by each mouse in the last session of the complex condition (c10). Using the targets A, B, and C as the Markov chain states would have provided a limited amount of data. Instead, we described states as movements to the left (L) or to the right (R) thereby obtaining larger pools of data (e.g. R = A → B, B →C, C → A) and a more compact description (e.g. two 0^th^ order groups instead of three). To assess the influence of rewards on mouse decisions when switching to the complexity condition (i.e. win-stay-lose-switch strategy), we also compared the probability of going forward P(F) or backward P(U) with the conditional probabilities given the presence or absence of reward (e.g. P(F_rw_) or P(U_unrw_)). In this case, F = R →R, L →L and U = R →L, R →L.

### Analysis of subsequences distribution

All patterns starting by A were extracted and pooled from the choice sequences of mice in the last sessions of the three conditions (training, complexity, probabilistic). The histograms represent the distribution of these patterns following the decision tree structure. In other words, two neighbor branches shared the same prefix.

### Bayesian model comparison

Bayesian model comparison aims to quantify the support for a model over another based on their respective likelihoods P(*D|M*), i.e. the probability that data D are produced under the assumption of model *M*. In our case, it is useful to compare the fitness of the *M*_*ind*_ model fitted session by session independently from that of the model *M*_*con*_ fitted to all sessions in a continuous way. Since these models do not produce explicit likelihood measures, we used approximate Bayesian computation: considering the 15 best fits (i.e. the 15 parameter sets that granted the highest fitness score), we estimated the models’ likelihood as the fraction of (*Ŝ, Ĉ* pairs that were within the confidence intervals of mouse data. Then, the Bayes factor was calculated as the ratio between the two competing likelihoods:

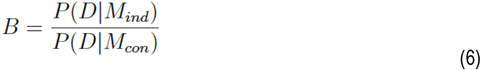

B > 3 was considered to be a substantial evidence in favor of *M*_*ind*_ over *M*_*con*_ ^30^.

### Statistical analysis

No statistical methods were used to predetermine sample sizes. Our sample sizes are comparable to many studies using similar techniques and animal models. The total number of observations (N) in each group as well as details about the statistical tests were reported in figure captions. Error bars indicate 95% confidence intervals. Parametric statistical tests were used when data followed a normal distribution (Shapiro test with p > 0.05) and non-parametric tests when they did not. As parametric tests, we used t-test when comparing two groups or ANOVA when more. Homogeneity of variances was checked preliminarily (Bartlett’s test with p > 0.05) and the unpaired t-tests were Welch-corrected if needed. As non-parametric tests, we used Mann-Whitney test when comparing two independent groups, Wilcoxon test when comparing two paired groups and Kruskal-Wallis test when comparing more than two groups. All statistical tests were applied using the scipy.stats Python module. They were all two-sided except Mann-Whitney. p > 0.05 was considered to be statistically non-significant.

## Acknowledgements

This work was supported by the Centre National de la Recherche Scientifique CNRS UMR 8246 et UMR 7222, the Labex SMART (ANR-11-LABX-65) supported by French state funds managed by the ANR within the Investissements d’Avenir programme under reference ANR-11-IDEX-0004-02, the Foundation for Medical Research (FRM, Equipe FRM DEQ2013326488 to P.F) and the French National Cancer Institute Grant TABAC- 16-022 (to P.F.). P.F. team is part of the École des Neurosciences de Paris Ile-de-France RTRA network and member of LabEx Bio-Psy.

## Author Contributions

PF designed the behavioral experiment. PF, EB, RDC, MD, ED, TAY, BH and MC performed the behavioral experiments, PF and MB analyzed the behavioral data. MB developed the computational model, SD developed some acquisition tools. JN and OS contributed to modeling studies and data analysis. MB, PF, JN and OS wrote the manuscript with inputs from AM.

## Supplementary Figures

**Supplementary Figure 1:**
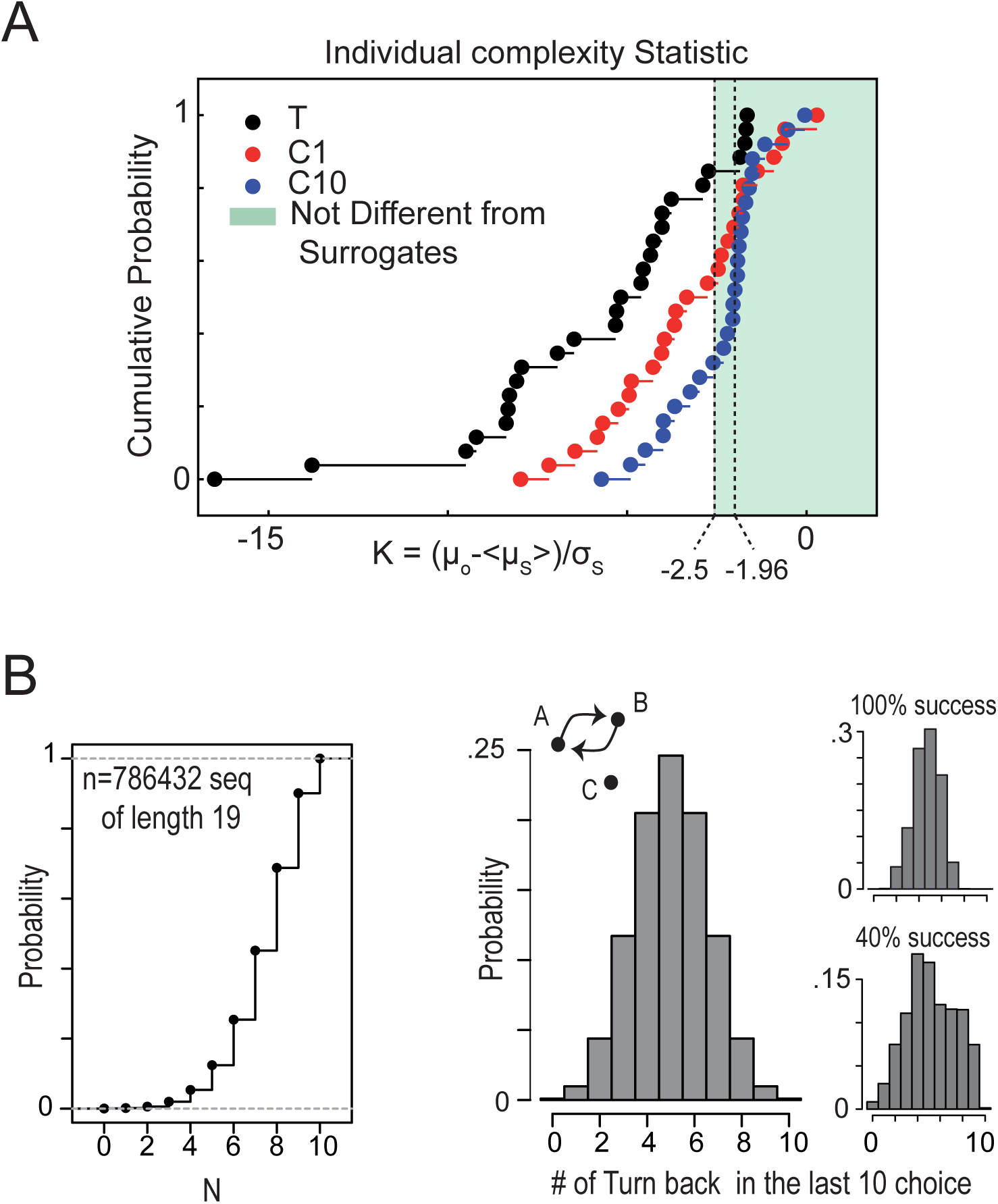
A) Individual comparison with random sequence of choices: Cumulative distribution of the K parameters calculated for each sequence of choices in sessions Training (T), C1 and C10. For each experimental sequence of choices, the term K=(µ_0_-<µ_s_>)/ σ_s_ was calculated, where µ_0_ is the complexity of the original data, <µ_s_>) and σ_s_ are the mean and standard deviation of the complexity of the surrogate series (i.e. for each experimental series, 1000 random sequences of the same length in which two consecutive elements could not be equal). We then tested the hypothesis that each original set was different from surrogates. Assuming Gaussian statistics, a limit of K=-2.5 and −1.96 indicates respectively a confidence of 99.4% and 95% that µ_0_ ≥ µ_s_. We found that 65.4% of mouse sequences in session C10 were not different from surrogates with a confidence of 99.4%. N = 27 in all sessions except C10 where N = 26. **B) Theoretical number of rewards and U-turns in the last 10 choices of a sequence of length 19:** *Left*: Cumulative distribution of the number of rewards obtained in the last 10 choices of the total number of sequences of length 19. Middle: Histogram of the number of U-turns in the last 10 choices. *Right*: Histogram of the number of U-turns in all possible sequences with 100% of reward (top) or 40% of reward (bottom). The optimal U-turn rate to maximize rewards is 50%.

**Supplementary Figure 2:**
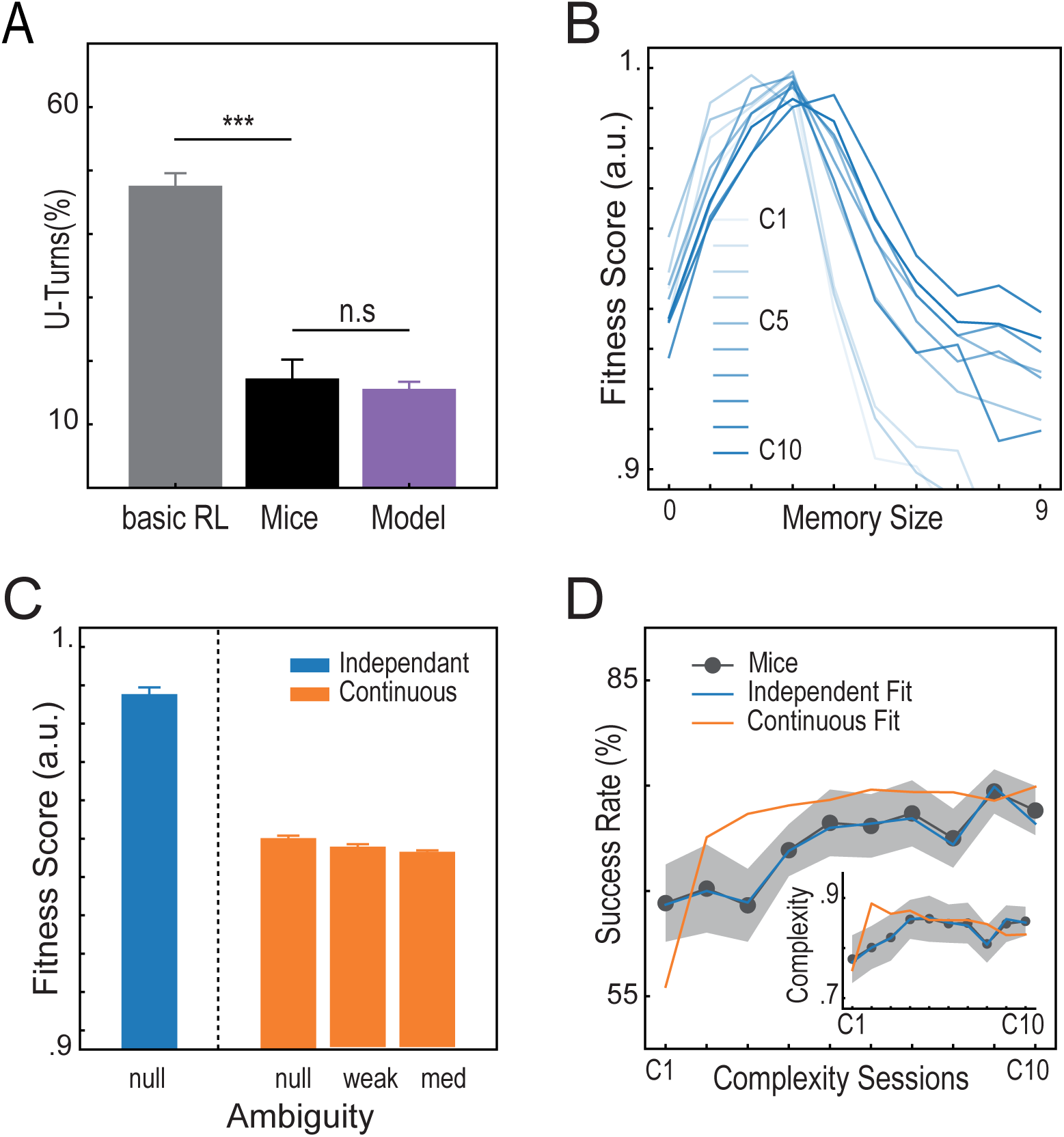
Model fitting results for different variants of the model. **A)** Comparison between the U-turn rates achieved by a basic RL algorithm, by our model and by mice in the training phase. A basic RL algorithm is unable to reproduce the stereotypical circular trajectories observed in mice during and after training (t(35) =−16.21, p = 10-17, Welch t-test; see Figure 1B for mouse trajectories). Indeed, with equal probabilities of reward at all targets, this algorithm learns equal state-action values and randomly chooses between them. Discounting the reward function by a U-turn cost representing previous locations in the state vector (see ‘Methods’) makes the model capable to generate the same percentage of U-turns as mice under the free condition (t(35) = 0.99, p = 0.32, Welch t-test). **B)**, **C)** and **D)** Comparison of fitness scores with independent and continuous model fitting procedures. ‘Independent’ refers to the session-by-session hyper parameter optimization, ‘continuous’ to the optimization of the model for all sessions in a row (see ‘Methods’). **B)** Best fitness scores obtained with each of the tested memory sizes session by session. Smaller memories fit better. **C)** Comparison of the fitness score obtained by the N = 15 best fits in the independent and continuous fitting procedure. In continuous fitting, three levels of state representation ambiguity are shown (see ‘Methods’). Error bars represent 95% confidence intervals. **D)** Success rates and complexity levels obtained in simulations in comparison with those obtained by mice. The shaded area represents the 95% confidence interval for mice. Only best fits are represented for model simulations (average of 20 runs). The model fitted for each session independently follows the evolution of behavioral data by adapting the exploration hyperparameter. By contrast, fitting all sessions with one single parameter set fails to reproduce the same evolution. In particular, the model exhibits a radical increase in success rate and complexity between sessions C01 and C02 as opposed to a more gradual progress in mice.

**Supplementary Figure 3:**
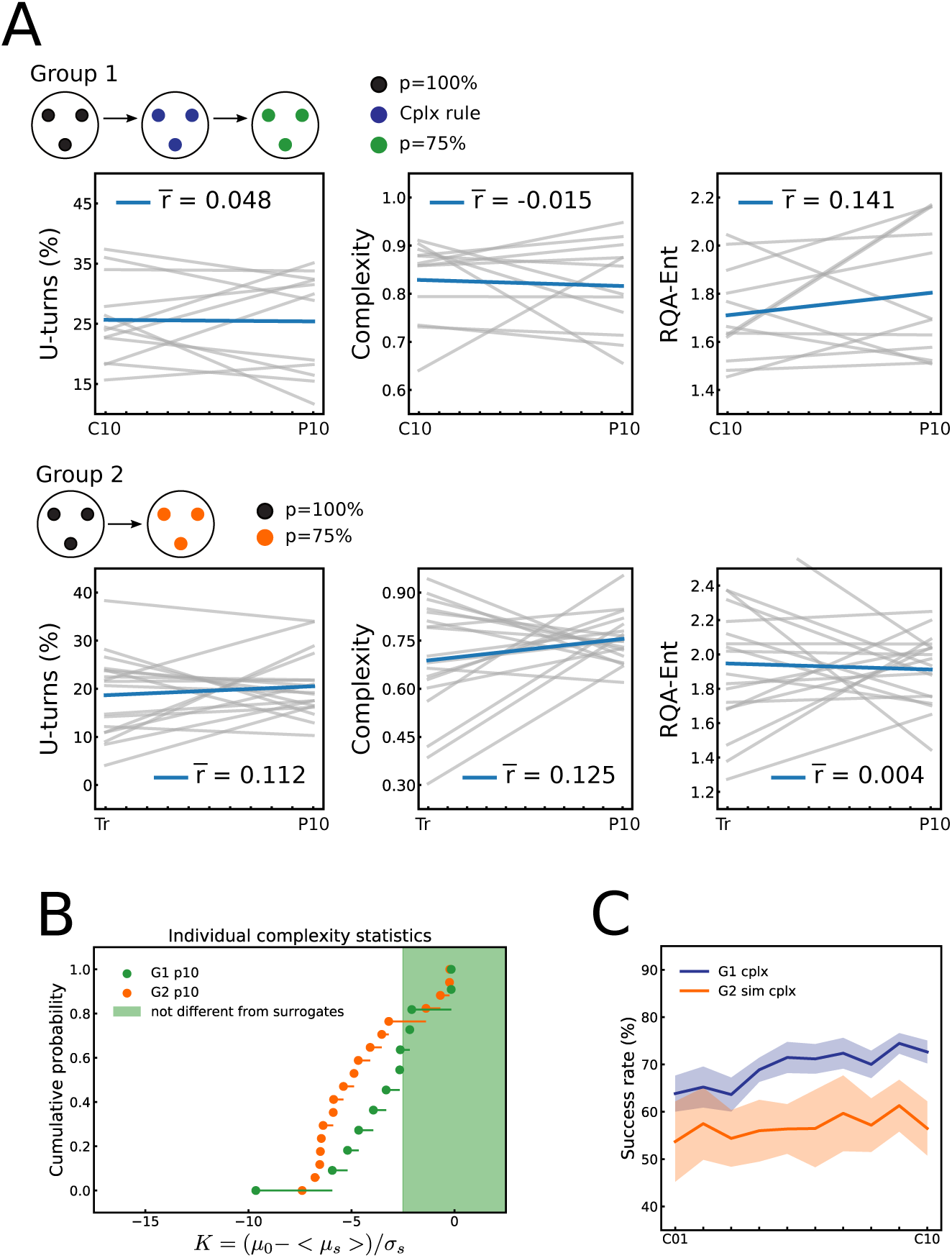
Absence of change in terms of U-turn rate, complexity and RQA entropy in the probabilistic condition compared to the previous condition. *Top*: Animals from Group 1. *Bottom*: Animals from Group 2. Grey lines represent linear regressions of the three measures for individual mice and the blue line represents the average. **B) Individual comparison with random sequence of choices:** Cumulative distribution of the K parameters calculated for each sequence of choices in the last sessions under the probabilistic rule p10 for G1 and G2 animals. For each experimental sequence of choices, the term K=(µ_0_-<µ_s_>)/ σ_s_ was calculated, where µ_0_ is the complexity of the original data, <µ_s_>) and σ_s_ are the mean and standard deviation of the complexity of the surrogate series (i.e. for each experimental series, 1000 random sequences of the same length in which two consecutive elements could not be equal). We then tested the hypothesis that each original set was different from surrogates. Assuming Gaussian statistics, a limit of K=-2.5 and −1.96 indicates respectively a confidence of 99.4% and 95% that µ_0_ ≥ µ_s_. We found that 33.3% of G1 mouse sequences and 22.3% of G2 mouse sequences were not different from surrogates with a confidence of 99.4%. N = 12 for G1, N = 18 for G2. **B) Success rate of the sequences performed by G2 animals if they were performed under the complexity rule:** We simulated the reward delivery rule of the complexity condition against the sequences performed by G2 mice under the probabilistic condition. Under these conditions, the animals would have reached a significantly lower success rate starting from the third session (G1 versus G2, session p01, U = 327, p = 0.053, Mann-Whitney test, session p02, t(28) = 1.81, p = 0.076, session p03, t(28) = 2.70, p = 0.0098, session p04, t(28) = 3.99, p = 0.0002, session p05, t(28) = 4.92, p = 0.00001, session p06, t(28) = 3.65, p = 0.00069, session p07, t(28) = 3.18, p = 0.0027, t-test, session p08, U = 394, p = 0.00049, Mann-Whitney test, session p09, t(28) = 4.85, p = 0.00001, session p10, t(28) = 5.55, p = 0.000002, t-test).

